# Plasma membrane rather than endosomal Gq signaling drives transcriptional activity by the viral chemokine receptor US28 in glioblastoma

**DOI:** 10.1101/2025.06.12.659276

**Authors:** Carole Daly, Adam Wright, Raimond Heukers, Chloe M. McKee, Rebecca C. Coll, Emma Evergren, Martine J. Smit, Alex R.B. Thomsen, Bianca Plouffe

## Abstract

US28 is a human cytomegalovirus-encoded chemokine receptor homologue that has high agonist-independent activity, internalizes constitutively, and plays an oncomodulatory role in glioblastoma. As G protein signaling was originally believed to strictly occur at the plasma membrane, it has been assumed that US28’s constitutive Gα_q/11_ signaling is mediated by a minor population at the plasma membrane. However, accumulating evidence shows that some GPCRs activate G proteins from intracellular organelles, such as endosomes. Importantly, endosomal rather than plasma membrane G protein signaling has been associated with transcriptional activity. Here, we demonstrate that the endosomal US28 population robustly activates Gα_q/11_, and thus, provides the major contribution of Gα_q/11_ signaling. Surprisingly, US28 signaling at the plasma membrane rather than from endosomes primarily drives upregulation of gene expression involved in cell proliferation and inflammatory responses that are associated with glioblastoma and cancer. Our findings highlight the crucial role of receptor signaling location in cellular responses.

## INTRODUCTION

Viral infections are estimated to be the direct cause of approximately 10-15% of human cancers worldwide^1^. Glioblastoma is the most aggressive and common primary brain tumor found in adults. Human cytomegalovirus (HCMV) has been detected in more than 84% of glioblastoma tumors and HCMV seropositivity correlates with glioblastoma progression as well as decreased life expectancy in patients^2, 3^. HCMV encodes four chemokine receptor homologues: UL33, UL78, US27 and US28^4^. Among these, US28, is the best characterized. This receptor displays robust constitutive activation of Gα_q/11_, which leads to cell proliferation, angiogenesis, inflammation, and metabolic reprogramming of tumor cells^5, 6, 7^. Notably, the constitutive activity of US28 increases cell proliferation of the host cell by stimulating the production and secretion of interleukin-6 (IL-6) and by enhancing sphingosine-1-phosphate receptor 1 (S1PR1) signaling, which both contribute to the activation of the signal transducer and activator of transcription 3 (STAT3) axis^8, 9^. An important STAT3 target gene that is upregulated in glioblastoma cells by US28/Gα_q/11_ activity is *CCND1* encoding cyclin D1, a protein involved in the G_1_ to S phase transition^9, 10^.

An interesting particularity of this receptor is its constitutive internalization via endocytic vesicles^11^. This constitutive internalization is due in part to agonist-independent phosphorylation of the carboxy-terminal of US28 by G protein-coupled receptor kinases (GRKs), subsequent recruitment of β-arrestins to the phosphorylated US28, and clathrin-dependent internalization^12, 13^. It has been estimated that ∼20 percent of the total receptor population is expressed at the plasma membrane, while the vast majority (∼80 percent) is located intracellularly^11^. Despite its robust agonist-independent activity, US28 binds several CC-type chemokines and CX3CL1 (fractalkine) with high affinity^14, 15, 16^. This property, combined with its constant internalization, result in chemokine scavenging by US28, which has been proposed as a strategy that HCMV-infected cells use to subvert and escape the host’s immune system^17^.

Like other chemokine receptors, US28 belongs to the superfamily of G protein-coupled receptors (GPCRs). Following the canonical model of GPCR signaling, extracellular agonist binds to the receptor at the plasma membrane, which leads to intracellular G protein engagement and activation. G protein activation initiates downstream signaling events that ultimately modulate cellular behavior. However, this initial signaling phase is short-lived. Recruitment of β-arrestins to the phosphorylated receptor uncouples it from G proteins and induces its internalization^18, 19^. These actions by β-arrestins were believed to terminate GPCR signaling. Therefore, for many years, GPCRs were thought to signal via G proteins exclusively at the plasma membrane. In line with this plasma membrane centric view of G protein signaling, it has been assumed that the constitutive Gα_q/11_ signaling by US28 was mediated solely by the minor population of receptors at plasma membrane^13^. However, the emergence of molecular tools to interrogate signaling events with a subcellular resolution provided several lines of evidence that certain GPCRs can continue to activate G proteins from endosomes after they have been internalized^20, 21, 22, 23, 24, 25^. Importantly, G protein signaling from endosomes by a specific receptor can initiate functionally distinct responses as compared to G protein signaling at plasma membrane by the same receptor, a concept that is termed ‘spatial bias’. One of the best characterized cellular responses that is regulated by endosomal GPCR signaling is gene transcription. Interestingly, US28 was recently reported to recruit sphingosine kinase 1 (SK1), an enzyme that directly interacts with activated Gα_q/11_ proteins, to early endosomes in a Gα_q_-dependent manner^9, 26^. Sphingosine-1-phosphate, produced by SK1 at endosomes, activates signaling pathways that influence gene transcription, thus connecting GPCR endosomal signaling to transcriptional regulation. This observation suggests the possibility that US28 activates Gα_q/11_ proteins from this intracellular compartment and regulates gene transcription.

G protein signaling from endosomes has been reported for several Gα_s_-coupled receptors such as parathyroid hormone receptor 1 (PTH1R)^27^, thyroid stimulated hormone receptor (TSHR)^28^, luteinizing hormone receptor (LHR)^29^, β_2_-adrenergic receptor (β_2_AR)^30^, vasopressin type 2 receptor (V_2_R)^24, 31, 32^, dopamine D1 receptor (D1R)^33^, glucose-dependent insulinotropic receptor (GIPR)^34^ and glucagon-like peptide-1 receptor (GLP-1R)^35^. For several of these receptors, inhibition of their internalization strongly diminishes agonist-mediated gene transcription. Using β_2_AR as a model, it has been demonstrated that cAMP-mediated transcriptional responses emanate from the endosome-localized receptor pool rather than receptor signaling at the plasma membrane^36, 37^. The current explanation for this spatial bias is that cAMP produced at the plasma membrane is contained locally. This is because phosphodiesterases hydrolyze cAMP within a narrow radius^38^, and therefore, cAMP never reaches the nuclear region when generated at the plasma membrane. In contrast, endosomes are located in close proximity to the nucleus and production of cAMP from these compartments triggers nuclear entry of cAMP-dependent protein kinase (PKA), where it phosphorylates cAMP-responsive element binding protein (CREB), a pivotal transcription factor^39^. Similarly to Gα_s_-coupled receptors, numerous GPCRs have been reported to activate Gα_q/11_ from endosomes, notably angiotensin type 1 receptor (AT_1_R)^40^, B2 bradykinin receptor (B2R)^40^, oxytocin receptor (OTR)^40^, thromboxane receptor (TPR)^40^, muscarinic acetylcholine receptor M3 (M3R)^40^, neurokinin 1 receptor (NK_1_R)^41, 42^, calcium-sensing receptor (CaSR)^43^, and protease-activated receptor-2 (PAR_2_)^44^. Whilst important cellular responses have been associated with endosomal Gα_q/11_ protein signaling, its role in regulating certain downstream responses and gene transcription in endogenous cellular systems is unclear. Moreover, as opposed to cAMP, the mechanisms underlying spatial regulation of the second messengers downstream of Gα_q/11_ activation (inositol 1,4,5-trisphosphate and diacylglycerol) have not been extensively studied. For many of the Gα_q/11_-coupled receptors, diacylglycerol has not yet been detected at endosomal membranes^40^, which complicates our understanding of the underlying signal transduction process that leads to cellular responses.

To characterize the subcellular localization of constitutively active US28 receptors and how gene transcription is potentially regulated in a spatially biased manner, we combined miniG protein (mG), nanobody technologies, mutagenesis, enhanced bystander bioluminescence resonance energy transfer (EbBRET), confocal microscopy, RNA sequencing, and physiological assays. Together, our data suggest that US28-mediated Gα_q/11_ signaling takes place at different endosomal subcellular compartments and that the location of this signaling plays a key role in directing oncomodulatory responses in glioblastoma cells.

## RESULTS

### US28 primarily activates Gα_q_ from endosomes

To define the subcellular location of active US28, we expressed mG proteins in human malignant glioblastoma astrocytoma tumor-derived U251 cells^45^. mG proteins are engineered GTPase domains of Gα subunits, which were initially developed for structural studies of active state GPCRs^46, 47^. mG proteins are localized in the cytosol, but are recruited to active GPCRs at the cell surface or endomembranes in a stable manner upon receptor stimulation. Several mG variants have been designed to capture the receptor coupling specificity of the four Gα subunit families (Gα_s_, Gα_i/o_, Gα_q/11_, Gα_12/13_)^47, 48^. To determine the subcellular location of active US28, we here used the mGsq_71 variant^47^, a mGs chimera with the specificity determinants of Gα_q_, referred hereafter as mGq for simplicity. For maximal sensitivity and robustness, we measured mGq recruitment to the plasma membrane and early endosomes using an EbBRET approach^49^. The energy acceptor (green fluorescent protein from *Renilla reniformis*; rGFP) was anchored at the plasma membrane or early endosomes by fusing its carboxy-terminus to the polybasic sequence and prenylation CAAX box of KRas^50^ (rGFP-CAAX) or Rab5^51^ (rGFP-Rab5), respectively (Fig. 1a, b). The energy donor, a highly luminescent mutant form of *Renilla reniformis* luciferase known as RlucII^52^ and referred hereafter as Rluc, was fused at its carboxy-terminal to mGq. Recruitment of mGq to rGFP-CAAX or rGFP-Rab5 increases the proximity between energy donor and acceptor, enabling resonance energy transfer from Rluc to rGFP, resulting in a rise in EbBRET values (Fig. 1a, b). Our data show that increasing US28 expression in U251 cells causes a small but significant increase of EbBRET values when rGFP-CAAX is used as energy acceptor compared to equivalent increasing expression of the inactive mutant US28-R129^3.50^A^53^ (Fig. 1a). The US28-R129^3.50^A mutant contains a mutated DRY motif which disrupts US28’s constitutive G protein activation and is considered an inactive US28 version. The modest activation at the plasma membrane is in line with the small population of wild-type US28 expressed at plasma membrane^11^. In contrast, the increase of EbBRET was much higher for similar amounts of US28 expressed when rGFP-Rab5 was used as energy acceptor, suggesting that early endosomes are rich in active US28 (Fig. 1b). As mG proteins are highly engineered and differ significantly from natural full-length Gα subunits, we also monitored the presence of active endogenous Gα_q/11_ at early endosomes. To do this, we expressed increasing amount of US28 using a previously developed EbBRET-based effector membrane translocation assay (EMTA)^40, 54^. This uses the sub-domain of p63RhoGEF, which selectively interacts with GTP-bound Gα_q/11_, fused at its carboxy-terminus to Rluc used as energy donor and rGFP-Rab5 as energy acceptor (Fig. 1c). Increasing amount of US28, but not US28-R129^3.50^A, induced a gradual recruitment of p63RhoGEF to Rab5 depicted by the progressive increase in EbBRET values (Fig. 1c). These results suggest that US28 robustly activates Gα_q/11_ at early endosomes (Fig. 1c).

**Fig. 1.**
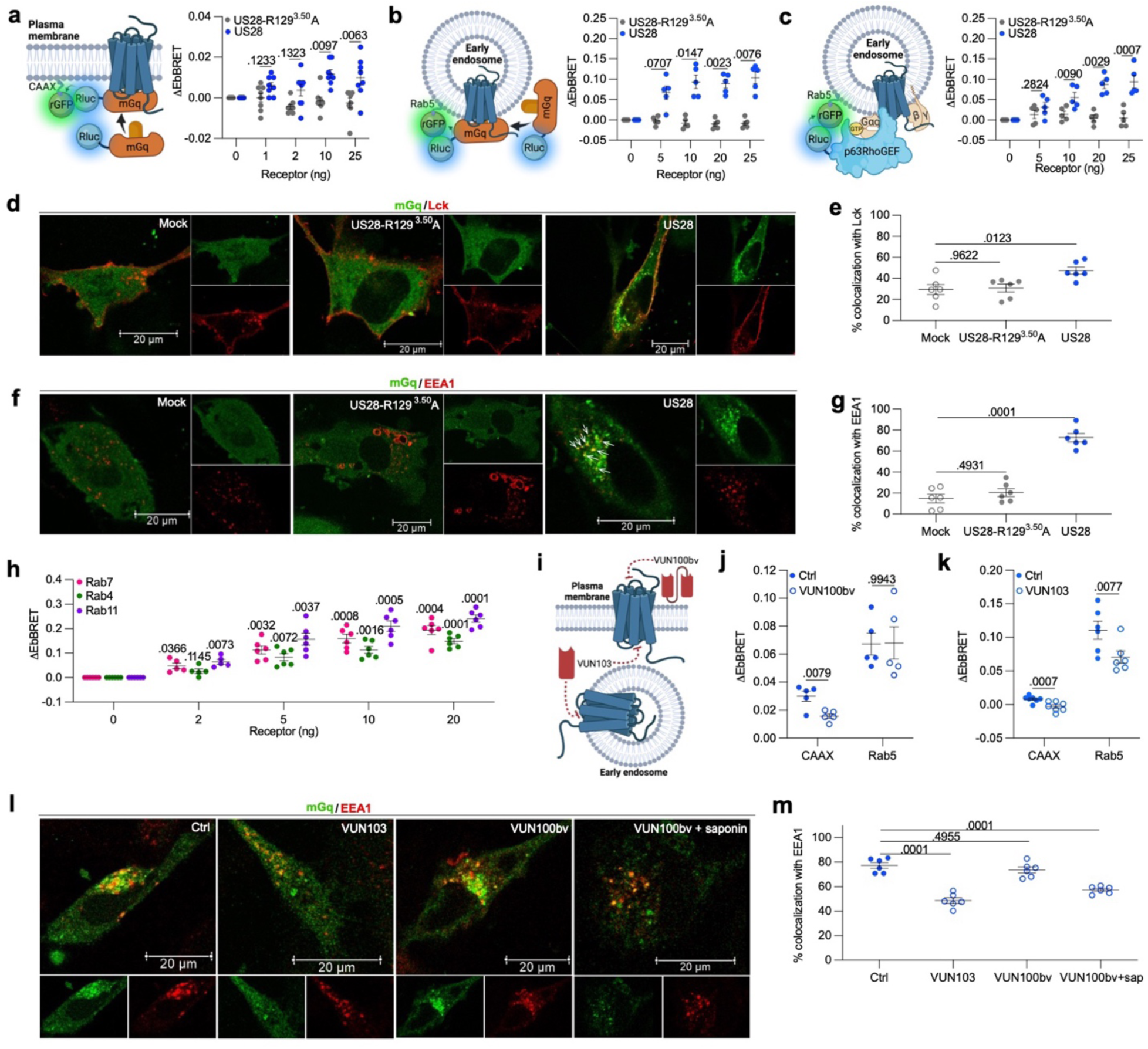
US28 primarily activates Gαq from endosomes in U251 cells. **a** Schematic illustration of the EbBRET assay monitoring energy transfer from mGq-Rluc to rGFP anchored at the plasma membrane by fusion with the CAAX box of KRas. Data represent US28-R129^3.50^A- or US28-mediated increase of EbBRET (*n* = 8). **b** Schematic representation of the EbBRET assay monitoring EbBRET between Rluc-mGq and early endosome-anchored rGFP-Rab5. Data represent US28-R129^3.50^A- or US28-mediated increase of EbBRET (*n* = 5). **c** Schematic illustration of the EbBRET assay monitoring EbBRET between Rluc-p63RhoGEF and rGFP-Rab5. Data represent US28-R129^3.50^A- or US28-mediated increase of EbBRET (*n* = 5). **d** Confocal microscopy (representative of 3 experiments) of cells expressing Halo-mGq, RFP-Lck (plasma membrane marker) and US28, US28-R129^3.50^A or empty vector (mock). **e** Percentage of mGq colocalized with Lck (from 6 representative images). **f** Confocal microscopy (representative of 3 experiments) of cells expressing Halo-mGq, RFP-EEA1 (early endosomes marker) and US28, US28-R129^3.50^A or vector alone (mock). **g** Percentage of mGq colocalized with EEA1 (from 6 representative images). **h** US28-mediated increase of EbBRET between Rluc-mGq and rGFP-Rab7, rGFP-Rab4, or rGFP-Rab11 (0, 5-20 ng; n = 6, 2 ng; *n* = 5). **i** Cartoon illustrating VUN100bv and VUN103 binding to the respective extracellular and intracellular regions of US28. **j** US28-mediated EbBRET between Rluc-mGq and rGFP-CAAX or rGFP-Rab5 in cells treated with 0.1 mM VUN100bv or vehicle (ctrl) (*n* = 5). **k** US28-mediated EbBRET between Rluc-mGq and rGFP-CAAX or rGFP-Rab5 in cells expressing VUN103 or empty vector (ctrl) (CAAX; *n* = 7, Rab5; *n* = 6). **l** Confocal microscopy (representative of 3 experiments) of cells permeabilized or not with saponin expressing Halo-mGq, US28, RFP-EEA1, VUN103 or empty vector (ctrl) and treated with 0.1 mM VUN100bv or vehicle (ctrl) (representative of 3 experiments). **m** Percentages of mGq colocalized with EEA1 in cells expressing VUN103 or empty vector treated with 0.1 mM VUN100bv or vehicle (ctrl) (from 6 representative images). All data represent means ± SEM. **a-c, j-k:** two-way ANOVA (Sidak’s post hoc test); **e, g, m:** one-way ANOVA (Dunnett’s post hoc test); **h:** two-way ANOVA (Dunnett’s post hoc test).

To further characterize the subcellular localization of US28 activity, we visualized U251 cells transfected with either a red fluorescent protein (RFP) fused to a plasma membrane (Lck^55^) or early endosome (EEA1; early endosome antigen 1^56^) marker along with Halo-tagged mGq (Halo-mGq) labeled with a cell permeable fluorescent HaloTag ligand using confocal microscopy. In addition to these markers, the cells were co-transfected with either US28, US28-R129^3.50^A or vector alone (mock). In mock and US28-R129^3.50^A transfected cells, mGq is homogenously distributed in the cytosol (Fig. 1d,f). In contrast, in presence of US28, mGq forms clusters in periphery of the nucleus depicting regions rich in receptors that couple to Gα_q/11_ (Fig. 1d,f). For cells expressing US28 and RFP-Lck, low levels of colocalization of mGq with Lck were observed (Fig. 1e). Although low (∼50%), the colocalization in presence of US28 was significantly higher than in cells expressing US28-R129^3.50^A or the vector alone (∼30%) (Fig. 1e). These data are therefore in line with our EbBRET data suggesting the presence of a minor population of active US28 receptors at the plasma membrane (Fig. 1a). In contrast, in cells expressing RFP-EEA1, ∼70% of EEA1 colocalized with mGq in presence of US28, which was significantly higher than RFP-EEA1-expressing cells transfected with US28-R129^3.50^A or vector alone (Fig. 1g). Interestingly, we observed that several mGq clusters did not colocalize with EEA1, which indicates that active US28 receptors may also be present in other types of endosomes (Fig. 1f). To investigate the nature of these other endosomes, we monitored US28-mediated EbBRET between Rluc-mGq and rGFP-fused Rab7^57^, Rab4^58^ and Rab11^59^, which are markers of late, fast recycling and slow recycling endosomes respectively (Fig. 1h). In agreement with our confocal microscopy observations, gradual increases of EbBRET signals were detected with all three energy acceptors when cells were transfected with increasing amounts of US28 (Fig. 1h). These data suggest that although early endosomes constitute a major site of active US28 receptors, late and recycling endosomes are also populated with active receptors.

To further validate that US28 receptors found at endosomes and plasma membrane are active, we took advantage of two inhibitory nanobodies targeting different regions of US28. The first nanobody (VUN100bv) is a high affinity bivalent nanobody that binds to the extracellular amino-terminal domain and loops of US28, and reduces US28-mediated constitutive accumulation of inositol phosphate and downstream signaling events^60^. The second high affinity nanobody (VUN103) binds to the second and third intracellular loops of US28 and displaces G protein. Thus, VUN103 inhibits constitutive US28-mediated accumulation of inositol phosphate^61^. We used purified VUN100bv as an extracellular ligand while VUN103 was expressed intracellularly as an ‘intrabody’ using plasmid DNA encoding VUN103. In whole-cell experiments, VUN100bv selectively inhibits the activity of the US28 population at the plasma membrane, since VUN100bv cannot cross the plasma membrane to access the intracellular receptors. In contrast, as an intrabody, VUN103 can access and inhibit both intracellular and plasma membrane receptor populations (Fig. 1i). We first measured the effect of VUN100bv on the population of active US28 receptors at the plasma membrane and early endosomes using our EbBRET-based mGq biosensors used in Fig. 1a and b, respectively. As sown in Fig. 1j, VUN100bv effectively blocks US28 activity at the plasma membrane, but not at early endosomes. These findings are in line with the inability of this nanobody to reach endosomal compartments. In contrast, VUN103 successfully blocked US28 activity at both the plasma membrane and early endosomes, in line with its intracellular location (Fig. 1k). To confirm that the inability of VUN100bv to reduce the endosomal population of active US28 receptors was linked to the extracellular location of this nanobody, we permeabilized our cells and used confocal microscopy to visualize the location of active receptors. Pretreatment with VUN100bv did not affect the recruitment of mGq to endosomes, unless cells were permeabilized with saponin, which caused significant VUN100bv-dependent inhibition (Fig. 1l,m). In contrast, in US28-expressing U251 cells, transfection of VUN103 significantly decreased the presence of mGq clusters and their colocalization with early endosomes (Fig. 1l,m). Altogether, our data indicate that despite a small population of active US28 receptors at the plasma membrane, the dominant active US28 receptor pool is located at endosomes.

### Mutation of the US28 carboxy-terminal phosphorylation sites relocates active receptors from endosomes to plasma membrane

To determine whether US28 activity at the plasma membrane and/or from endosomes regulates gene expression, we sought to shift the main pool of US28 from the endosomes to the plasma membrane. As the carboxy-terminus of US28 is phosphorylated in an agonist-independent manner, β-arrestins are constitutively associated with US28, which leads to steady-state internalization of the receptor^13^. To reduce US28 internalization and thereby increase its expression at the cell surface, we used a mutant US28 receptor harboring a **<u>P</u>**hospho**<u>D</u>**eficient **C**arboxy-**T**erminus for which all serine and threonine residues of the carboxy-terminus were respectively replaced by alanine and valine and referred to as US28-PDCT (Fig. 2a).

**Fig. 2.**
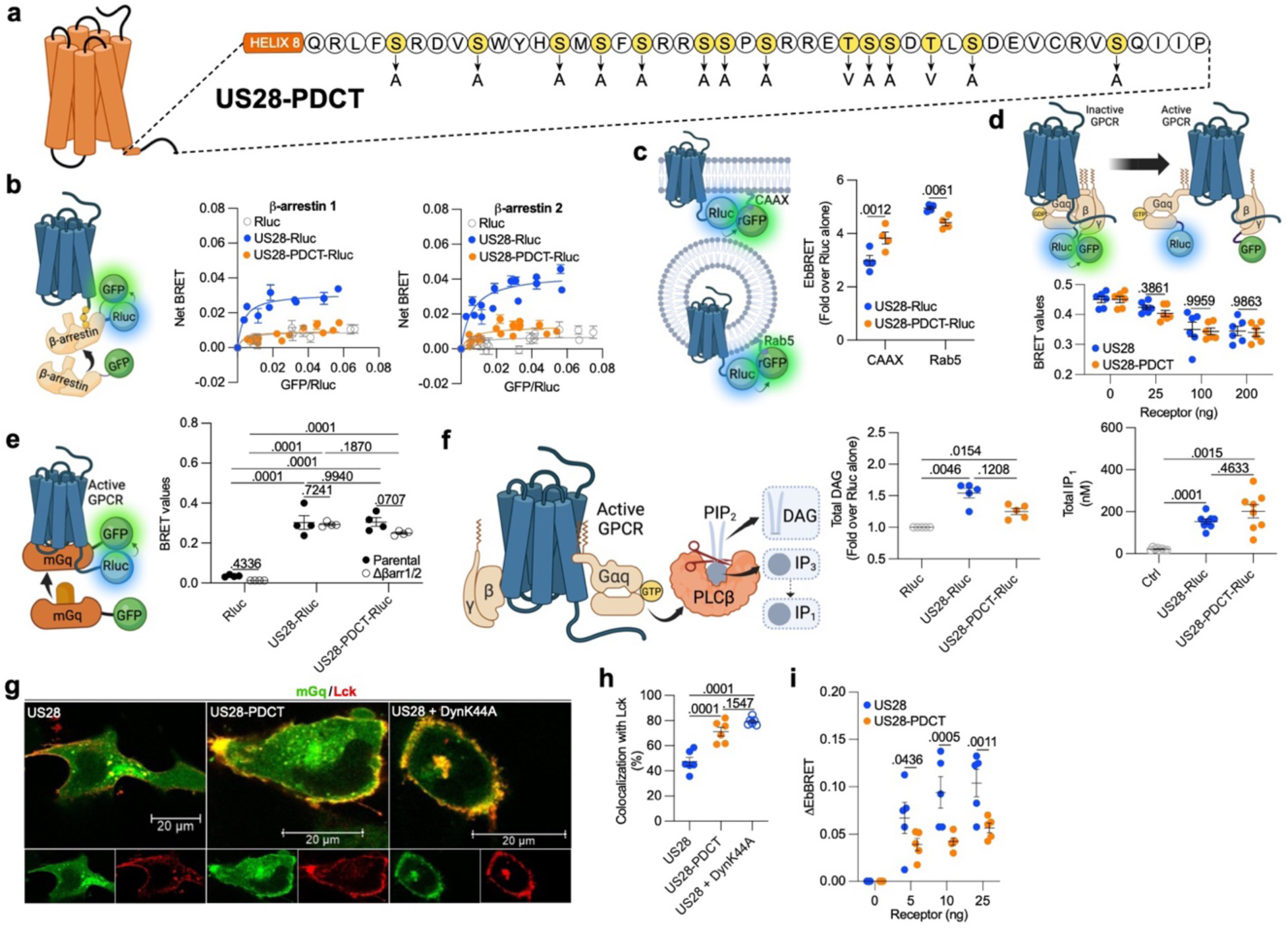
Characterization of a mutant US28 harboring a phospho-deficient carboxy-terminal (US28-PDCT) in U251 cells. **a** Cartoon showing US28-PDCT with serine (S) and threonine (T) residues after helix 8 mutated to alanine (A) or valine (V). **b** Cartoon of the BRET assay monitoring interaction between β-arrestins-GFP and US28-Rluc or US28-PDCT-Rluc. Data represent net BRET titration curves obtained by increasing GFP-β-arrestins 1/2 with fixed amounts of US28-Rluc/US28-PDCT-Rluc/Rluc. Each point represents the mean ± SEM of net BRET values measured in quadruplicates. The points from each experiment (β-arrestin 1; *n* = 3, β-arrestin 2; *n* = 4) were pooled and fitted using a non-linear one site specific binding relationship. **c** EbBRET assay monitoring US28-Rluc and US28-PDCT-Rluc expression at plasma membrane and early endosomes by recording EbBRET between US28-Rluc or US28-PDCT-Rluc and rGFP-CAAX or rGFP-Rab5. Data represent EbBRET values between Rluc-fused receptors and rGFP-CAAX or rGFP-Rab5 over EbBRET values obtained with Rluc (*n* = 4). See Supplementary Fig. 1. **d** BRET assay monitoring energy exchange from Rluc-Gα_q_ to GFP-Gγ_1_. Data represent the BRET values with increasing receptors (*n* = 6). **e** BRET assay to compare the active conformations of US28 and US28-PDCT, which monitors mGq-GFP recruitment to US28-Rluc, US28-PDCT-Rluc or Rluc. Data show BRET values with US28-Rluc, US28-PDCT-Rluc or Rluc and mGq-GFP in β-arrestin-deficient or parental HEK-293 cells (*n* = 4). See Supplementary Fig. 2. **f** Cartoon representing second messengers downstream of receptors. Data represent relative DAG levels in cells expressing US28-Rluc, US28-PDCT-Rluc or Rluc and IP_1_ concentrations in cells expressing US28-Rluc or US28-PDCT-Rluc (DAG; *n* = 5, IP_1_; *n* = 8). See Supplementary Fig. 3. **g** Confocal images of cells expressing Halo-mGq, RFP-Lck along with US28, US28- PDCT without/with DynK44A (representative from 3 experiments). **h** Percentage of Halo-mGq/RFP-Lck colocalization in cells expressing US28, US28-PDCT with/without DynK44A (from 6 representative images). **i** Relative EbBRET values in cells expressing rGFP-Rab5/Rluc-mGq and increasing levels of US28 or US28-PDCT (*n* = 5). All data represent means ± SEM. **c-d, i:** two-way ANOVA (Sidak’s post hoc test), **e**: two-way ANOVA (Tukey’s post hoc test), **f, h**: one-way ANOVA (Tukey’s post hoc test).

To confirm the loss of β-arrestin recruitment to US28-PDCT, we used a BRET-based approach where Rluc is fused to the carboxy-terminus of US28-PDCT and US28, respectively, while GFP10^62^ was fused to β-arrestins 1 and 2 (Fig. 2b). In this configuration, the constitutive recruitment of β-arrestins to the receptors generates a BRET signal. Our BRET saturation curves confirm that US28 constitutively interacts with β-arrestin 1 and β-arrestin 2 (Fig. 2b) as BRET values are higher with US28-Rluc than with Rluc alone (background BRET). In contrast, the BRET values with US28-PDCT-Rluc are undistinguishable from the BRET values obtained with Rluc alone, reflecting a loss of constitutive interaction with β-arrestins for US28-PDCT (Fig. 2b). To verify that this loss of interaction with β-arrestins translates to increased receptor expression at the plasma membrane and a corresponding reduction of receptors at endosomes, we measured EbBRET between Rluc-fused receptors and rGFP-CAAX or rGFP-Rab5 (Fig. 2c). At similar total expression levels (Supplementary Fig. 1), our results show a significant increase of US28-PDCT expression at the plasma membrane and a decrease at the early endosomes as compared to US28 (Fig. 2c). We also verified that the ability of US28-PDCT to constitutively activate Gα_q/11_ was not changed by the loss of receptor interaction with β-arrestins. This was confirmed by first measuring the gradual dissociation of Gα_q_ from Gγ_1_, as an index of Gα_q/11_ activation (Fig. 2d). As the constitutive activation monitored is proportional to the number of receptors present, increasing expression of US28 or US28-PDCT resulted in progressive decrease of BRET between Rluc-fused Gα_q_ and GFP10-fused Gγ_1_ (Fig. 2d). As no statistical difference was detected between US28 and US28-PDCT for the same amount of receptors transfected, our results suggest an equivalent ability of US28-PDCT and US28 to constitutively activate Gα_q/11_ (Fig. 2d). To further demonstrate that the loss of β-arrestin binding to US28-PDCT does not increase the constitutive activation of Gα_q/11_, we monitored the recruitment of mGq fused to GFP10 directly to US28-Rluc and/or US28-PDCT-Rluc by BRET in parental and CRISPR/Cas9-engineered β-arrestin 1/2-deficient HEK293 cells (Fig. 2e). At similar levels of receptor expression (Supplementary Fig. 2), no difference in mGq recruitment to US28 and US28-PDCT was detected (Fig. 2e). Furthermore, the absence of β-arrestins in cells expressing US28 or US28-PDCT did not increase mGq recruitment to these receptors, suggesting that the loss of interaction between β-arrestins and US28-PDCT does not increase the constitutive activity of this receptor (Fig. 2e). We also verified that the total constitutive production of diacylglycerol (DAG) and inositol-3-phosphate (IP_3_) resulting from the hydrolyzation of phosphatidylinositol 4,5-bisphosphate (PIP_2_) by phospholipase Cβ (PLCβ) activated downstream of Gα_q/11_ were similar for US28 and US28-PDCT (Fig. 2f). Our results indicate that at equivalent total expression of US28-Rluc and US28-PDCT-Rluc (Supplementary Fig.3), no statistical difference was detected between US28- and US28-PDCT-mediated production of DAG and inositol monophosphate (IP_1_ – a metabolite of IP_3_) (Fig. 2f), in line with the similar ability of US28 and US28-PDCT to constitutively activate Gα_q/11_.

To verify that the US28-PDCT rescue of expression at the plasma membrane translates to a higher number of active receptors at the plasma membrane, we used the same confocal microscopy approach as in Figure 1d. We observed that expression of US28-PDCT by U251 cells led to increased colocalization of mGq with the plasma membrane marker Lck as compared to cells expressing wild-type US28 (Fig. 2g,h). Moreover, similar enhanced levels of Lck colocalized mGq was observed for wild-type US28 upon co-expression of the dominant-negative dynamin DynK44A, which interferes with receptor internalization^63^ (Fig. 2g,h). As there are more active receptors at the plasma membrane in case of US28-PDCT compared to US28, we hypothesized that conversely less active US28-PDCT should be present in endosomes. Our data indeed confirmed that mGq recruitment to early endosomes was lower for US28-PDCT as compared to US28 and independent of receptor expression (Fig 2i). Altogether, our data confirm that the displacement of US28 from endosomes to the plasma membrane also relocates the subcellular location of Gα_q/11_ signaling without changing the intrinsic constitutive coupling to Gα_q/11_.

### US28 relocation to the plasma membrane increases the number of upregulated cancer- and cell division-associated genes

Endosomal signaling is known to regulate transcriptional activity by several GPCRs via Gα_s_ or Gα_q/11_. Therefore, we hypothesized that the constitutive US28-mediated Gα_q/11_ signaling from endosomes, rather than at the plasma membrane, drives transcription of oncomodulatory genes.

To investigate this, we transfected U251 cells with either US28-Rluc, US28-PDCT-Rluc or Rluc alone. The luminescence emitted by Rluc was monitored to ensure equal total receptor expressions (Supplementary Fig. 4). Total RNA from these cells was extracted and sequenced (Supplementary Table S1). Two differential gene expression analyses (DGEAs) were performed to determine differentially regulated genes in U251 cells expressing US28-Rluc or US28-PDCT-Rluc versus Rluc alone (Supplementary Table S2-3, Fig. 3a). We found that 330 or 594 genes were upregulated in U251 cells expressing either US28-Rluc or US28-PDCT-Rluc, respectively (Fig. 3b, Supplementary Table S2-3), compared to Rluc-expressing U251 cells. From these genes, the expression of 247 was enhanced by both US28-Rluc and US28-PDCT-Rluc (Fig. 3b, Supplementary Table S4). Thus, out of the 330 genes upregulated by US28-Rluc, only 83 were not upregulated by US28-PDCT-Rluc (Fig. 3b-c, Supplementary Table S4). Interestingly, 347 genes were only upregulated by US28-PDCT-Rluc (Fig. 3b and d, Supplementary Table S4). These results suggest that in contrast to our hypothesis, US28 located at the plasma membrane is a stronger regulator of gene expression compared to endosomally-located US28. Additionally, the transcription of several genes does not seem to depend on the subcellular location of US28.

**Fig. 3.**
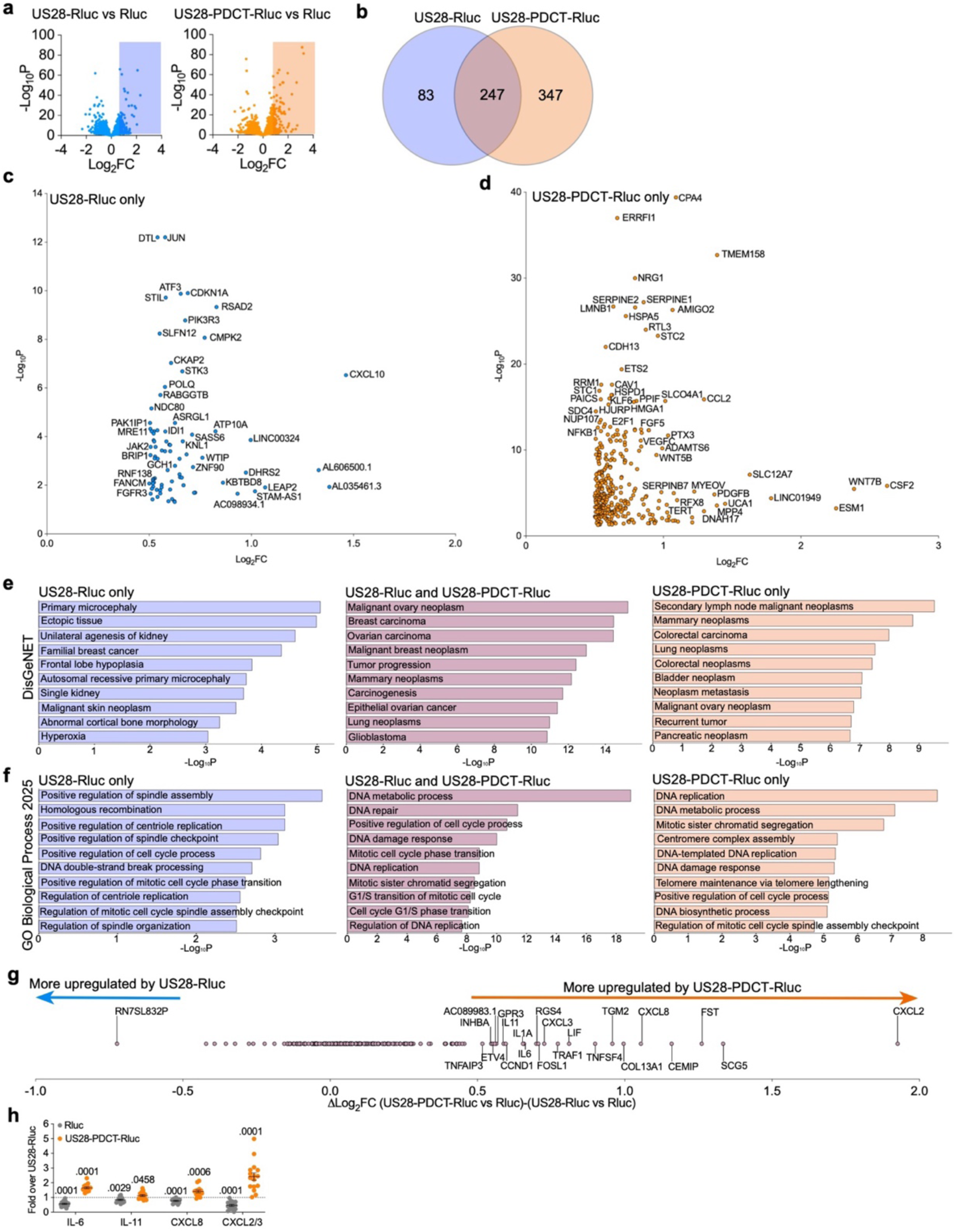
US28 signalling at plasma membrane drives upregulation of genes associated to cancer and cell division. **a** Volcano plot representing the genes regulated by US28-Rluc and US28-PDCT-Rluc in U251 cells compared to Rluc alone (*n* = 3). See also Supplementary Fig. 4, and Supplementary Tables 2-3. **b** Venn diagram representing the genes upregulated by US28-Rluc only, by US28-PDCT-Rluc only, and by both receptors. See Supplementary Table 4. **c** Volcano plot of the 83 genes upregulated by US28-Rluc only. **d** Volcano plot of the 347 genes upregulated by US28-PDCT-Rluc only. **e** Bar graphs representing the top-10 enriched terms in DisGeNET library with associated p values for the genes upregulated by US28-Rluc only, by both receptors, and by US28-PDCT-Rluc only. See also Supplementary Fig. 5. **f** Bar graphs representing the top-10 enriched terms in GO Biological Process 2025 library with associated p values for genes upregulated by US28-Rluc only, by both receptors, and by US28-PDCT-Rluc only. See also Supplementary Fig. 6. **g** Difference of Log_2_FC values between (US28-PDCT-Rluc vs Rluc) and (US28-Rluc vs Rluc) for the 247 upregulated by US28-Rluc and US28-PDCT-Rluc. **h** Scatter plot representing the relative concentration of cytokines secreted in the conditioned media by U251 cells expressing US28-Rluc, US28-PDCT-Rluc, or Rluc alone (*n* = 15 for IL-6 and IL-11, *n* = 14 for CXCL8, *n* = 16 for CXCL2/3). Data are expressed as mean ± SEM and *p* values (one sample t test) versus US28-Rluc (set to 1) are shown. See also Supplementary Fig. 7.

A pathway analysis by Enrichr^64, 65, 66^ was conducted where each gene set was used as input data in DisGeNET, a database of human gene-disease associations^67^. Our search showed a strong correlation between the upregulated genes and various types of cancers, including glioblastoma (Fig. 3e, Supplementary Fig. 5). This correlation is particularly strong for genes that are upregulated by both US28-Rluc and US28-PDCT-Rluc (Fig. 3e, Supplementary Fig. 5). Furthermore, using the GO Biological Process 2025 database^68, 69^, we found that many of the upregulated genes are tied to cellular processes that are dysfunctional in cancers, including DNA replication/metabolism and cell cycle regulators (Fig. 3f, Supplementary Fig. 6). Again, this correlation is more significant for genes that are enhanced by both US28 and US28-PDCT (Fig. 3f, Supplementary Fig. 6).

Among the most interesting genes upregulated by both US28-Rluc and US28-PDCT-Ruc, but to a greater extent by US28-PDCT-Ruc, are *CCND1* and *IL6,* which have been reported to increase glioblastoma cell proliferation^8, 10^. Interestingly, several other genes encoding for cytokines and chemokines, such as *IL11*, *IL1A*, *LIF*, *TNFSF4*, *CXCL3*, and *CXCL8* were also upregulated by both US28-Rluc and US28-PDCT-Ruc, but to a greater extent by US28-PDCT-Ruc (Fig. 3g). These data suggest that, in addition to driving the upregulation of more genes, US28-PDCT-Rluc also enhances expression of certain genes that are upregulated by US28-Rluc, but to a greater extent.

To investigate whether the increased cytokine and chemokine transcription in US28-PDCT-expressing U251 cells also leads to an increase in their secretion, we measured IL-6, IL-11, CXCL8, CXCL2, and CXCL3 concentrations in conditioned media. In agreement with our DGEA (Supplementary Table S2), the relative levels of IL-6, IL-11, CXCL8, CXCL2, and CXCL3 present in the media of cells expressing US28-Rluc were higher than in the media of cells expressing Rluc alone (Fig. 3h, Supplementary Fig. 7). This indicates that expression of US28 in glioblastoma cells increases the secretion of these cytokines and chemokines. Importantly, their relative levels in the media of cells expressing US28-PDCT-Rluc were even higher than in the media of cells expressing equivalent amount of US28-Rluc (Fig. 3h). Altogether, our data implies that US28 signaling primarily at the plasma membrane drives the upregulation of genes associated with cancer and cell cycling. The increased expression of genes encoding cytokines and chemokines further translates into their translation and secretion, which potentially modifies the tumor micro-environment.

### US28 at the plasma membrane enhances US28-mediated growth of glioblastoma spheroids and neutrophil chemotaxis

US28-mediated upregulation of cyclin D1 and IL-6 secretion are key factors that promote proliferation of the host cells^8, 10^. The genes encoding these proteins are more upregulated in and IL-6 more secreted from glioblastoma cells expressing US28-PDCT-Rluc compared to US28-Rluc expressing cells (Fig. 3g,h). Furthermore, US28-PDCT-Rluc upregulates several additional genes also linked to cell cycle (Fig. 3f, Supplementary Fig. 6). Therefore, we hypothesized that US28 activity at the plasma membrane may be more potent at inducing cell proliferation than US28 activity at endosomes. To test this hypothesis, we generated U251 cells stably expressing equivalent levels of US28-Rluc or US28-PDCT-Rluc (Supplementary Fig. 8a). Next, U251 cell spheroids were grown from a clone expressing US28-Rluc and two clones expressing US28-PDCT-Rluc. Spheroids offer a 3D model that better replicate cell growth of solid tumors compared to 2D cell cultures. Their circular area was measured each day for a period of 10 days (Fig. 4a-c). Additionally, the number of cells forming the spheroids was determined at day 10 (Fig. 4d). Supporting our hypothesis, the spheroids expressing US28-PDCT-Rluc showed a greater proliferation than the spheroids expressing US28-Rluc, depicted by their higher circular area (Fig. 4a-c) and a greater increase in the total number of cells from the spheroids (Fig. 4d) at day 10. There was no statistical difference between the rate of proliferation of spheroids between the two clones expressing US28-PDCT-Rluc (Fig. 4b-d). These results suggest that Gα_q/11_ signaling by US28 at the plasma membrane is superior to US28 signaling at endosomes in promoting cell proliferation.

**Fig. 4.**
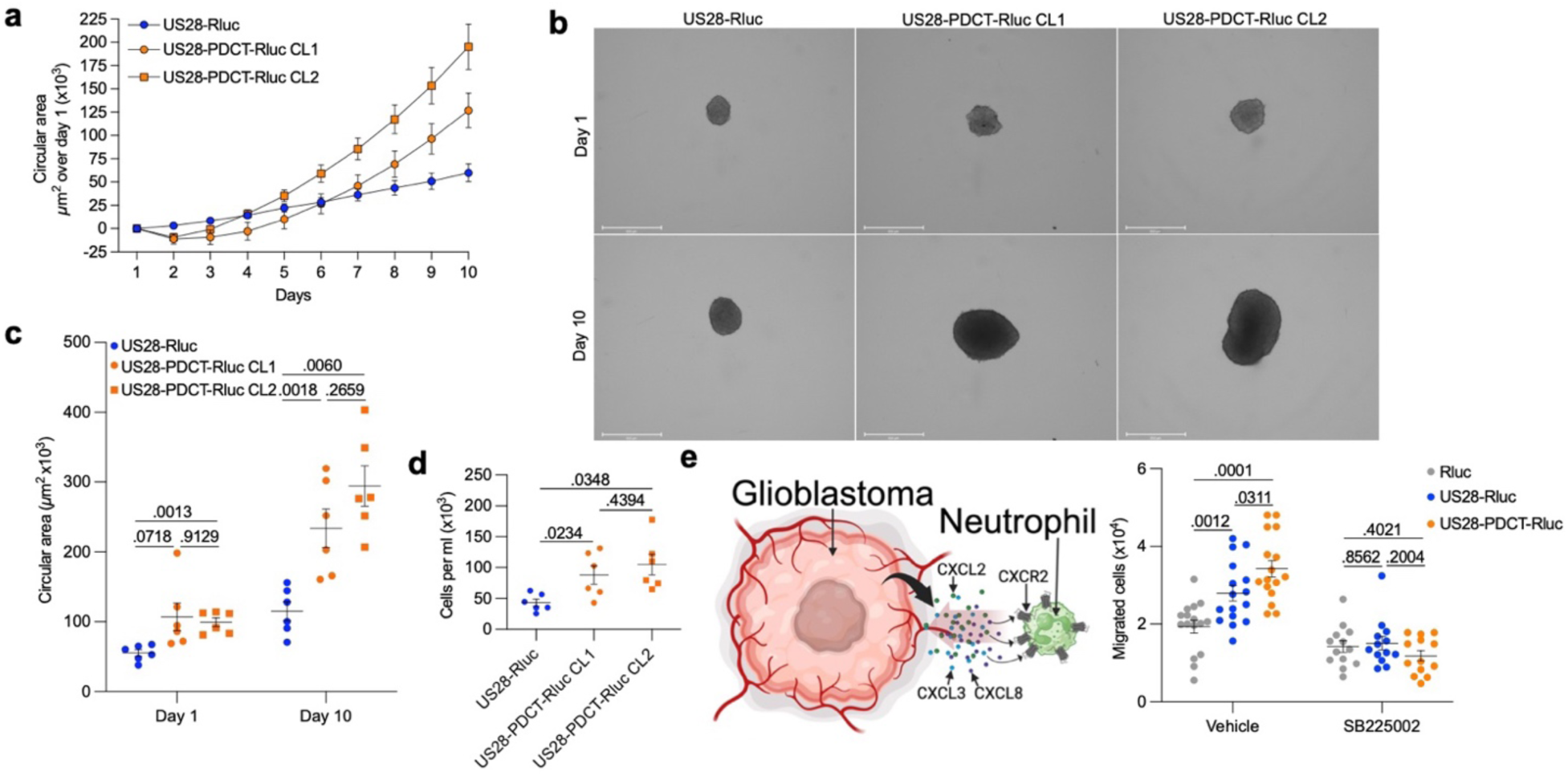
Relocation of active US28 from endosomes to plasma membrane enhances US28-mediated growth of glioblastoma spheroids and neutrophil chemotaxis. **a** Growth curve of glioblastoma spheroids stably expressing US28-Rluc or US28-PDCT-Rluc (*n* = 6). 2 different clones (CL1 and CL2) for US28-PDCT-Rluc. See also Supplementary Fig. 8a. Data represent the net increase in circular area over day 1. **b** Representative images of glioblastoma spheroids on day 1 and day 10 stably expressing US28-Rluc or US28-PDCT-Rluc (*n* = 6). **c** Scatter plot representing the circular area of glioblastoma spheroids stably expressing US28-Rluc or US28-PDCT-Rluc measured at day 1 and day 10 (*n* = 6). **d** Number of cells per ml obtained by dissociation of the cells composing the glioblastoma spheroids stably expressing US28-Rluc or US28-PDCT-Rluc on day 10 (*n* = 6). **e** Number of differentiated HL-60 cells preincubated with the CXCR2 inhibitor SB225002 or vehicle migrated toward the conditioned media from U251 cells expressing US28-Rluc or US28-PDCT-Rluc (vehicle; *n* = 16, SB225002; *n* = 13). See also Supplementary Figure 8b. All data represent means ± SEM. **c, e:** two-way ANOVA (Tukey’s post hoc test), **d**: one-way ANOVA (Tukey’s post hoc test).

To further validate the importance of cell surface Gα_q/11_ signaling by US28 in the upregulation of oncomodulatory genes and associated cellular responses, we focused on the increased expression and secretion of CXCL2, CXCL3, and CXCL8 from U251 cells expressing US28-PDCT-Rluc (Fig. 3g-h). As CXCL2, CXCL3 and CXCL8 are agonists of the chemokine receptor CXCR2^70^, we probed whether conditioned media collected from U251 cells expressing either US28-Rluc, US28-PDCT-Rluc or Rluc alone could provoke CXCR2-mediated responses. Mature circulating neutrophils express high levels of CXCR2, which directs their migration to sites of inflammation^71^. Therefore, we hypothesized that the conditioned media collected from U251 cells expressing US28-PDCT-Rluc would attract more neutrophils compared to the conditioned media from U251 cells expressing US28-Rluc or Rluc alone. To test this hypothesis, chemotaxis was monitored in a transwell assay using differentiated neutrophil-like HL-60 cells, which are derived from lymphocytes from a patient with acute promyelocytic leukemia. The migration of differentiated HL-60 cells towards a chamber containing conditioned media from US28-Rluc, US28-PDCT-Rluc, or Rluc expressing U251 cells was assessed (Supplementary Fig. 8b, Fig. 4e). In line with the higher levels of CXCL2, CXCL3 and CXCL8 secreted into the media of U251 cells expressing US28-Rluc compared to Rluc alone (Fig. 3h), more differentiated HL-60 cells migrated towards the media collected from these cells (Fig. 4e). Moreover, in agreement with our hypothesis and the higher concentrations of neutrophil chemoattractants found in the media collected from US28-PDCT-Rluc-expressing U251 cells (Fig. 3h), we also observed increased chemotaxis of differentiated HL-60 cells towards media from US28-PDCT-Rluc-expressing U251 cells relative to media from US28-Rluc-expressing cells (Fig. 4e). Importantly, the levels of chemotaxis were CXCR2-dependent as preincubation of the differentiated HL-60 cells with the CXCR2 antagonist SB225002^72^ abolished the chemotaxic response (Fig. 4e).

Altogether, we have demonstrated that the upregulation of oncomodulatory genes by US28 signaling at the plasma membrane triggers stronger cellular responses of the host cells contributing to the oncomodulatory nature of US28.

## DISCUSSION

US28 is widely expressed in human glioblastoma and displays oncomodulatory activity by activating a broad panel of transcription factors, such as NF-kB, nuclear factor of activated T cell (NFAT), CREB, STAT3, activator protein-2 (AP-2), stimulating protein 1 (Sp1), TCF/LEF, and hypoxia inducible factor-1 (HIF-1)^7, 8, 10, 73, 74, 75, 76^. A wide array of genes is under transcriptional control of these factors and our RNA sequencing data provided a comprehensive transcriptional profile of the genes upregulated by US28 in a glioblastoma context. In addition to previously reported *IL6* and *CCND1*, we identified 368 additional genes that are robustly upregulated by US28 (Fig. 3b). Importantly, many of these genes are associated with glioblastoma, (Fig. 3e, Supplementary Fig. 5), which highlights the oncomodulatory role of this viral GPCR in brain cancer. Interestingly, several other genes are related to other types of cancers, such as breast, ovary, and lung (Fig. 3e, Supplementary Fig. 5), which implies that US28 also enhances more general oncomodulatory pathways. This is exemplified by several genes involved in DNA processes and positive regulation of cell cycle (Fig. 3f, Supplementary Fig. 6). Further investigation is required to define the respective role of these individual genes in the development of glioblastoma, but our results suggest that US28 promotes robust transcriptional activity via constitutive activation of Gα_q/11_. As US28 is trafficked into intracellular compartments in a ligand-independent fashion, US28 is an ideal model to study the role of Gα_q/11_ location bias on gene transcription.

Applying molecular tools to monitor the subcellular location of receptors and Gα proteins in their active states, such as mG proteins and EMTA, confirmed our hypothesis that not only the minor population of US28 at the plasma membrane constitutively activates Gα_q/11_, but that the major endosomal receptor pool also does so. Previous studies using mutants of US28 with truncated carboxy-terminal or US28^ST/A^ reported contradicting data. In these studies, similar or higher levels of constitutive production of total inositol phosphate and activation of NF-kB were reported for these mutants^12, 13, 53^. However, the total expression of each mutant relative to wild-type was not assessed, which may potentially explain the differences observed in these publications. In the present work, the subcellular location of signaling was well defined and the respective total expression of US28 and US28-PDCT were matched using a luminescent tag to eliminate any misleading result caused by differences in expression rather than biological effect.

Several GPCRs have been reported to activate Gα_q/11_ from early endosomes and important cellular responses are regulated by this endosomal Gα_q/11_ signaling. For example, the release of substance P from primary sensory neurons following painful stimulus activates NK_1_R in second-order spinal neurons and triggers NK_1_R internalization into endosomes. Importantly, Gα_q/11_ signaling by internalized NK_1_R is responsible for the sustained activation of spinal neurons and pain transmission^41, 42^. In line with the key role of the endosomal compartment as a NK_1_R signaling hub that mediates sustained nociception, targeting NK_1_R antagonists to endosomes by conjugation to cholestanol or by modifications to increase their retention in this compartment, provides potent and long-lasting pain relief in a mouse model^41, 42^. In a heterologous cellular model that overexpresses NK_1_R, the reduction of NK1R internalization decreases the activation of the serum response element transcription factor mediated by substance P, suggesting a role of endosomal signaling by NK_1_R in transcriptional activity^41^. However, as NK_1_R also activates Gα_s_/cAMP signaling in a receptor internalization-dependent manner^41^, it is unclear if the transcriptional activity is mediated by endosomal Gα_s_ or Gα_q/11_ signaling. PAR_2_ is another receptor that mediates nociception via activation of Gα_q/11_ from endosomes^44^. This compartmentalized signaling is reported to promote pain sensation in patients with irritable bowel syndrome (IBS)^44^. Colonic mucosa from IBS patients releases proteases that induce PAR_2_ endocytosis, endosomal signaling, and persistent hyperexcitability of nociceptors. Accumulation of PAR_2_ antagonists in endosomes by conjugation to cholestanol or inhibition of trypsin-induced PAR_2_ internalization was shown to suppress persistent hyperexcitability and allodynia in vivo. In a heterologous cellular system, stimulation of PAR_2_ with trypsin induced colocalization of PAR_2_ with Gα_q/11_ at early endosomes, accumulation of inositol phosphate and activation of ERK at the nucleus. Importantly, PAR_2_ internalization is required for the activation of ERK at the nucleus and blocking Gα_q/11_ impaired nuclear ERK activation. Although these findings suggest a potential role of endosomal Gα_q/11_ signaling on transcriptional activity, gene transcription was not monitored downstream of nuclear ERK activation and the role of endosomal Gα_q/11_ signaling by internalized PAR_2_ has not been verified in vivo.

CaSR regulates extracellular calcium by endocrine actions in parathyroid glands, intestine, kidney, and bones. In these tissues, CaSR couples to Gα_q/11_, Gα_i/o_, and Gα_12/13_^77^. Patients with familial hypocalciuric hypercalcemia type-3, a disorder due to mutations of the adaptor protein-2 (AP2) leading to defective clathrin-mediated endocytosis, have an increased expression of CaSR at the plasma membrane^78^. As this condition is associated with reduced CaSR signaling via Gα_q/11_, a subsequent study demonstrated that CaSR preferentially signals via Gα_q/11_ from endosomes^43^. In that study, the impaired CaSR internalization led to reduced levels of inositol phosphate production, extracellular calcium-induced intracellular calcium release, ERK activation, and membrane ruffling. Most importantly, reduced CaSR internalization was also associated with a decrease in activation of serum responsive element (SRE), suggesting that endosomal Gα_q/11_ signaling may mediate gene transcription^43^. However, the authors also noticed CaSR-mediated changes in cAMP production which were partially dependent of Gα_q/11_. Therefore, this system is not ideal in dissecting the contribution of endosomal Gα_q/11_ signaling on transcriptional activity. Although there is a clear role of endosomal Gα_q/11_ signaling on cellular responses and physiology, its contribution to gene transcription is not as obvious as that of Gα_s_-coupled receptors.

Novel molecular tools with subcellular resolution recently developed have confirmed the presence of active Gα_q/11_ and colocalization of GPCRs coupled to Gα_q/11_ in active conformations at early endosomes^40, 41, 42, 43, 44^. However, the mechanisms of signal transduction downstream of Gα_q/11_ activation from endosomes remain unknown. The possible presence of PLCβ at endosomes has not been investigated yet and while phosphatidylinositol 4,5-bisphosphate (PIP_2_), the canonical substrate of this enzyme, is abundant at plasma membrane, it is almost absent from endosomal membranes. Therefore, an unanswered question is how does Gα_q/11_ activation from endosomes translate to a cellular response? Our data demonstrate that although US28 constitutively activates Gα_q/11_ at the plasma membrane and from endosomes, Gα_q/11_ signaling at plasma membrane seems to drive a more robust transcriptional response, which is in line with the efficacy of VUN100bv to impair US28-enhanced GBM growth^60^. Additionally, our data suggest that the transcription of some genes is independent from US28 subcellular location. More studies are required to determine if US28 is unique, or rather represents a proof-of-concept that endosomal Gα_q/11_ signaling, as opposed to Gα_s_ signaling from endosomes, does not result in a transcriptional response as robust as at the plasma membrane.

## METHODS

### Chemical reagents and antibodies

Coelenterazine 400a was purchased from NanoLight Technology. Linear 25 kDa polyethyleneimine (PEI) was supplied by Polysciences. HaloTag^®^ Oregon Green^®^ ligand was from Promega. Saponin solution 0.1% in PBS, 4% paraformaldehyde in PBS, Dulbecco’s modified Eagle’s medium (DMEM), Roswell Park Memorial Institute (RPMI) 1640 medium, Dulbecco’s phosphate-buffered saline (DPBS) and penicillin-streptomycin were purchased from ThermoFisher. The bivalent anti-US28 nanobody VUN100bv has been previously detailed^60^. The anti-human diacylglycerol rabbit polyclonal antibody conjugated to HRP was obtained from LSBio (cat# LS-C710932-LSP). YM-254890 was supplied by Tocris. All other chemicals and compounds were from Sigma Aldrich.

### Plasmids

US28 and US28-R129^3.50^A tagged with Flag at their N-termini in pTwistCMV expression vector were synthetized from Twist Bioscience. Gβ_1_ in pcDNA3.1+ was from www.cdna.org. C-tRFP-Lck cloned into PCMV6-AC-RFP expression vector was purchased from Origene (#RC100049). TagRFP-T-EEA1 in pEGFP-C1 vector and K44A HA-dynamin 1 in pcDNA3.1 were respective gifts from Silvia Corvera and Sandra Schmid (Addgene plasmids #42635 and #34683). β-arrestin1-GFP10 and rGFP-Rab7 in pcDNA3.1+ was a kind gift from Prof. Michel Bouvier (Université de Montréal, Canada) and Stéphane Laporte (McGill University, Canada). *Renilla reniformis luciferase* II (Rluc), US28-Rluc, and US28-PDT-Rluc in pcDNA3.1+ were synthetized by GenScript. Rluc-mGq and Rluc-mGs in pTwistCMV were synthetized by Twist Bioscience. The Venus tag in NES-Venus-mGsq and Venus-mGs previously described^48^ were replaced by Rluc. Halo-tagged mGq was kindly provided by Prof. Nevin A. Lambert (Augusta University, USA). The plasmid encoding for the intrabody VUN103 has been previously detailed^61^. All other plasmids used in this study have been previously described: rGFP-CAAX^49^, rGFP-Rab5^49^, rGFP-Rab7, rGFP-Rab4^49^, rGFP-Rab11^49^, p63-RhoGEF-Rluc^54^, βarrestin2-GFP10^79^, Gα_q_-118-RlucII (referred as Rluc-Gα_q_)^80^, GFP10-Gγ_181_.

### Cell Culture and transfection

U251 MG (human malignant glioblastoma astrocytoma) cells (Sigma Aldrich) were used for all experiments unless stated otherwise. These cells were trypsinized in trypsin-EDTA 0.05% and cultured in DMEM supplemented with 10% fetal bovine serum, penicillin (100 units per ml), and streptomycin (100 mg per ml), maintained at 37°C and 5% CO_2_ and passaged every 5-6 days. Transfections were performed 24 hours after seeding cells in 96-well plates (10,000 cells/well). A total of 1 µg DNA was combined with linear 25 kDa PEI at a ratio of 4 µg PEI per µg of DNA in DPBS for U251 cells or 3 µg PEI per µg of DNA in DPBS for HEK293 cells. Salmon sperm DNA (Thermo Fisher) or pcDNA3.1+ empty vector (Invitrogen) was used to complete to 1 µg total DNA. DNA mixed with PEI (100 µl total) was incubated for at least 15 minutes before distributing 10 µl per well (10,000 cells). Cells were incubated at 37°C, 5% CO_2_ for 48 hours before assay.

### Bioluminescence Resonance Energy Transfer (BRET)-based assays

Cells were seeded in white 96-well plates (Greiner) at 10,000 cells/well. Transfected cells were washed with DPBS and assayed in Tyrode’s buffer [137 mM NaCl, 0.9 mM KCl, 1 mM MgCl_2_, 11.9 mM NaHCO_3_, 3.6 mM NaH_2_PO_4_, 25 mM Hepes, 5.5 mM glucose, 1 mM CaCl_2_ (pH 7.4)] at 37°C. Nanobodies or vehicle were added and cells incubated at 37°C for the required time. 5 minutes before reading, 2.5 µM of coelenterazine 400a was added. All BRET measurements were performed using a FLUOstar Omega microplate reader (BMG Labtech) with an acceptor filter (515 ± 30 nm) and donor filter (410 ± 80 nm). BRET was calculated by dividing GFP emission by Rluc emission. The Rluc emission (RLU) from the donor filter was used to determine the total expression of receptor fused to Rluc.

### Confocal microscopy

Cells were seeded in 8-well glass chambered slides (Ibidi GMBH) at 30,000 cells/well, transfected the next day and assayed 48 hours upon transfection. To label Halo-mGq, HaloTag® Oregon Green® ligand was added to cells at a final concentration of 1 µM in the media and incubated 15 minutes (37°C, 5% CO_2_). Cells were washed 3 times with the media and incubated 30 minutes for the last wash (37°C, 5% CO_2_). The media was replaced by Tyrode’s buffer. To permeabilize the cells, 0.1% saponin in PBS was added and incubated for 10 minutes with the cells (37°C, 5% CO_2_). Cells were washed 3 times with DPBS, incubated 10 minutes for the last wash and DPBS was replaced by Tyrode’s buffer. Cells were stimulated with VUN100bv or vehicle for the required time at 37°C, 5% CO_2_. The media was removed and cells were fixed with 300 µl per well of 4% paraformaldehyde in PBS and incubated at room temperature for 10 minutes. The paraformaldehyde solution was replaced by DPBS and cells were incubated for 10 minutes before being replaced by 300 µl of Tyrode’s buffer and visualized on a SP8 confocal microscope (Leica) at 63X magnification. Images were quantified using Imaris cell imaging software version 9.9.1 (Bitplane, Oxford instruments). For each image, the surface module was selected isolating cells containing the region of interest. Thresholds for each channel were established be selecting the lower intensity from the region of interest (plasma membrane or endosomes). Data was reported as red volume (red voxels) above the threshold that is colocalized with green volume (green voxels) above the threshold and expressed as percentage.

### ELISA to monitor total cellular diacylglycerol

For the ELISA, 50 μl per well of a poly-D-lysine solution (Cultrex; 0.1 mg per ml) was added and the plates incubated at 37 °C for at least 30 min. Following the incubation, the poly-D-lysine solution was aspirated and wells washed twice with DPBS before adding 100 μl per well of a U251 cell suspension (100,000 cells per ml). Four additional wells in a separate white 96-well plate were seeded to monitor the RLU (receptor expression). The next day, cells were transfected as described. 48 hours after transfection, the RLU was monitored on the white plate as described in the BRET-based assays section. The media from the ELISA plate was removed, cells were washed with DPBS and fixed with 4% paraformaldehyde in PBS (50 μl per well) during 10 minutes at room temperature followed by three washes with 100 μl per well of washing buffer (0.5% BSA, 0.1% Triton-X100 in DPBS). Cells were incubated 10 min at room temperature for the last wash. The buffer was replaced with 50 μl per well of HRP-conjugated diacylglycerol antibody (1:1000 in washing buffer) and cells were incubated 1 hour at room temperature. Cells were washed three times with the washing buffer and three times with DPBS. To detect HRP, cells were incubated with 100 μl per well of SigmaFast^TM^ OPD solution as recommended by the manufacturer and the reaction was stopped by adding 25 μl per well of HCl 3M. 100 μl was transferred into a transparent 96-well plate and the absorbance was monitored at 492 nm using a FLUOstar Omega microplate reader.

### IP_1_ assay

Parental U251 cells or U251 cells stably expressing US28-Rluc or US28-PDT-Rluc were seeded (100 μl; 80,000 cells per well) in quadruplicate in two white 96-well plates and incubated overnight at 37°C, 5% CO_2_. The next day, one plate was used to monitor the luminescence from Rluc (as an index of total receptor expression) as described in BRET-based assays section. The other plate was used to determine the concentration of IP_1_ using a CisBio IPOne ELISA kit according to the manufacturer procedures. Briefly, the media was aspirated and 30 μl per well of stimulation buffer (10 mM Hepes, 1 mM CaCl_2_, 0.5 mM MgCl_2_, 4.2 mM KCl, 146 mM NaCl, 5.5 mM glucose, 50 mM LiCl pH 7.4) was added. The plate was incubated 60 minutes (37°C, 5% CO_2_) and 30 μl of 1% lysis buffer was added to each well followed by 30 minutes incubation (37°C, 5% CO_2_). The lysate from the four wells from a same condition was combined and 50 μl of sample (or standards) was seeded on the ELISA plate provided in the kit. In each well, 25 μl of IP_1_-HRP conjugate and 25 μl of anti-IP_1_ monoclonal antibody (Mab) were added immediately. The plate was sealed and incubated on a plate shaker at 200 rpm for 3 hours at room temperature. The wells were washed six times with 250 μl of 1x washing buffer per well. The buffer was aspirated, replaced by 100 μl per well of the HRP substrate (TMB) and the plate was incubated in the dark at room temperature for 20-30 minutes before adding 100 μl of stop solution per well. The absorbance was monitored at 450 and 620 nm using a FLUOstar Omega microplate reader. The values read at 620 nm (optical imperfections in plate) were subtracted from the values read at 450 nm. The final IP_1_ concentration in the samples were determined using a range (0 – 5 μM final) of IP_1_ standard solutions derived from the calibrator provided in the kit.

### mRNA sequencing

Cells were seeded in 60 mm petri dishes (500,000 cells per dish) and transfected with US28, vector alone, US28-Rluc, or US28-PDT-Rluc in triplicate. 48 hours upon transfection, luminescence from Rluc was measured to ensure total expression was similar between conditions. Total RNA was extracted using a PureLink^TM^ RNA Mini Kit (Invitrogen) per the manufacturer’s instructions. Fragment analysis was performed and libraries were generated using a KAPA Stranded RNA HyperPrep kit with Riboerase HMR, normalized and pooled. The balanced pool was run on a NextSeq 2000 (Illumina) flow cell at >20M paired end reads per sample.

### Differential Gene Expression Analysis (DGEA)

To identify the genes statistically upregulated by US28-Rluc and US28-PDT-Rluc, DGEAs were performed by comparing US28-Rluc to Rluc alone samples and US28-PDT-Rluc to Rluc alone from the FASTQ files generated. Log_2_ fold change (Log_2_FC) along with -log_10_P (P referring to adjusted *P* value) values were calculated for US28-Rluc vs Rluc and for US28-PDT-Rluc vs Rluc comparisons genes. Upregulation was considered significant if -log_10_P > 1.3 and Log_2_FC > 0.5. From 60,583 genes analyzed, 677 genes met these statistical criteria.

### Quantification of cytokines and chemokines secreted

The quantification of IL-6, IL-11, CXCL8, CXCL2, and CXCL3 in the cell culture supernatant of U251 cells expressing US28-Rluc, US28-PDT-Rluc, or Rluc was performed using sandwich ELISA kits from Raybiotech detecting the human proteins according to the manufacturer procedures. Briefly, 48 hours after transfection, the cell media was removed and frozen at -80°C until assayed. The cells were washed with DPBS and replaced by Tyrode’s buffer. The luminescence from Rluc was measured to ensure similar expression of the receptors (or Rluc alone) as described in BRET-based assays section. The cell supernatant was diluted in cell growth media (1/50 for IL-6, 1/2 for IL-11, 1/1000 for CXCL8, and 1/10 for CXCL2/3). 100 µl of culture media, diluted supernatant or standard was added to each well of a 96-well plate coated with antibodies and incubated 2.5 hours at room temperature. 100 µl of prepared biotin antibody was added to each well and incubated 1 hour at room temperature. 100 µl of HRP-conjugated streptavidin solution was added to each well and incubated 45 minutes at room temperature. 100 µl per well of HRP substrate (3,3,5,5’-tetramethylbenzidine solution) was added and the wells were incubated 30 minutes at room temperature. The reactions were stopped by addition of 50 µl per well of 0.2 M sulfuric acid solution and the absorbance read at 450 nm using a FLUOstar Omega microplate reader. The concentrations of cytokines and chemokines from the growth media were below the level of detectability and therefore not considered.

### Generation of spheroids to monitor U251 proliferation

A 3 ml cell suspension (10,000 cells per ml) was prepared with U251 cells stably expressing US28-Rluc as well as with 2 clones of U251 cells expressing US28-PDT-Rluc (CL1 and CL2). The cell suspensions were centrifuged at 5,000 rpm for 5 minutes. The media over the cell pellets was replaced by 3 ml of phenol red-free media and the cells from each pellet were gently put back in suspension by trituration. 100 µl of each of these cell suspensions was seeded in 4 wells of a white 96-well plate and coelenterazine 400a was added as described in BRET-based assays to monitor the RLU emitted by Rluc (total receptor expression). 200 µl of the remaining cell suspensions were seeded per well in a clear and sterile 96-well CellStar® U-bottom plate with cell-repellent surface (Greiner). The plate was sealed, centrifuged at 1250 rpm for 5 minutes and put in a 37°C, 5% CO_2_ incubator for 24 hours before carefully removing 100 µl of media and replacing by 100 µl of fresh media. The spheroids were imaged each day over 10 days using a EVOS FL Auto 2 microscope (Invitrogen) at 40x magnification. The measure of the circular area was performed using ImageJ (Fiji package) using the freehand selection tool to outline the perimeter of each spheroid. The last day, the cells from each spheroid were resuspended using trypsin-EDTA 0.5% and counted.

### Neutrophil migration assay

HL-60 were differentiated to neutrophils by incubation in RPMI media containing 0.5% FBS, 2% nutridoma, 1.3% DMSO, and 1% penicillin-streptomycin for 5 to 7 days. 200,000 differentiated cells were suspended in 100 µl culture medium and transferred into Costar® Transwell® polycarbonate membrane inserts with 5 µm pore size (Millipore Sigma). Each insert was fitted into a well in a 24 well plate containing 500 µl of conditioned culture media. Cells were allowed to migrate from the top insert to the lower well chamber of the plate for 30 min at 37 °C at 5% CO_2_. At the end of the experiment, all migrated cells from the lower chamber were collected and transferred to a Countess™ Cell Counting Chamber Slide (Thermo Fisher Scientific) and counted using a Countess™ 3 Automated Cell Counter (Thermo Fisher Scientific). Cells used for assessment of CXCR2 involvement, were pretreated with 10 µM CXCR2 antagonist (SB225002) or vehicle DMSO control for 30 min prior to the experiment.

### Data and statistical analysis

In all experiments at least three independent experiments were performed and for each experiment *n* value is provided in the corresponding figure legend. GaphPad Prism version 10.4.1 was used to analyze all data. Statistical significance was calculated using post hoc multiple comparisons tests (Dunnet’s, Tukey’s or Sidak’s) after one-way or two-way ANOVA analysis or using one sample t test when data compared to normalized values (0 or 1), as indicated in respective figure legends. Summarized data are presented on graphs as mean ± SEM, and exact *p* values are provided on graphs.

## DATA AVAILABILITY

The data that support this study are available from the corresponding authors upon request. All data generated and analyzed during this study are included in this published article and the Supplementary Information. Source data are provided with this paper.

## ACKNOWLEDGEMENTS

This study was supported by a Wellcome Trust Seed Award (215229/Z/19) and a New Investigator Award from the UK Research and Innovation (UKRI) Biotechnology and Biological Sciences Research Council (BBSRC) (BB/X002578/1) to B.P. and by the LEO Foundation (LF18043), and NIH grants R35GM147088 (NIGMS) and R21CA243052 (NCI) to A.R.B.T. M.J.S. and R.H. were supported by the Dutch Research Council (NOW: Vici grant 016.140.657). R.C.C. was supported by a Research Grant from the UKRI Medical Research Council (MR/Y014065/1) and a New Investigator Award from the UKRI BBSRC (BB/V016741/1). C.D. and A.W. were supported by a Postgraduate Studentship from the Northern Ireland Department for the Economy (DfE). We thank Nick Bergkamp for producing VUN100bv nanobody. The cartoons were generated with BioRender.com.

## AUTHOR CONTRIBUTIONS

B.P. and A.R.B.T. take responsibility for the data integrity and accuracy of data analysis. B.P. and A.R.B.T. conceived and designed the study. C.D., A.W., C.M.M., A.R.B.T., and B.P. performed experiments and analyzed the data. R.H. and M.J.S. provided reagents. E.E., R.C.C., M.J.S., A.R.B.T, and B.P. coordinated, supervised and oversaw the study. C.D. drafted the initial version of the manuscript with support from B.P. B.P. and A.R.B.T. wrote the final version of the manuscript with input from all authors.

