## Supplementary information for "Plasma membrane rather than endosomal Gq signaling drives transcriptional activity by the viral chemokine receptor US28 in glioblastoma"

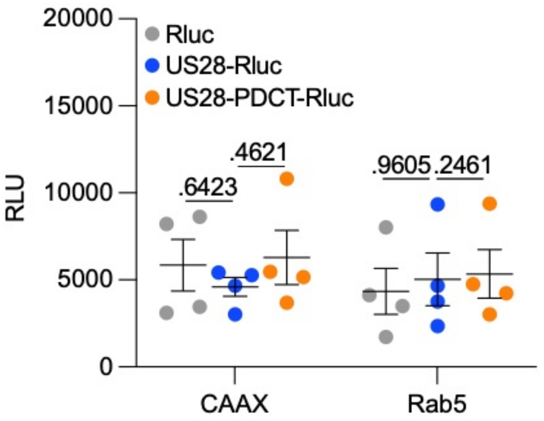


**Supplementary Fig. 1: Similar total expression of US28-Rluc, US28-PDCT-Rluc, or Rluc alone across U251 cells transfected transiently to evaluate the subcellular expression of these constructs at the plasma membrane (CAAX) or early endosomes (Rab5).** Data represent the luminescence expressed as relative luminescence units (RLU) monitored in U251 cells transiently transfected with US28-Rluc, US28-PDCT-Rluc or Rluc alone. Data are shown as means ± SEM of *n* = 4 independent biological replicates performed in technical quadruplicates and were analyzed by two-way ANOVA with Sidak's post hoc test.

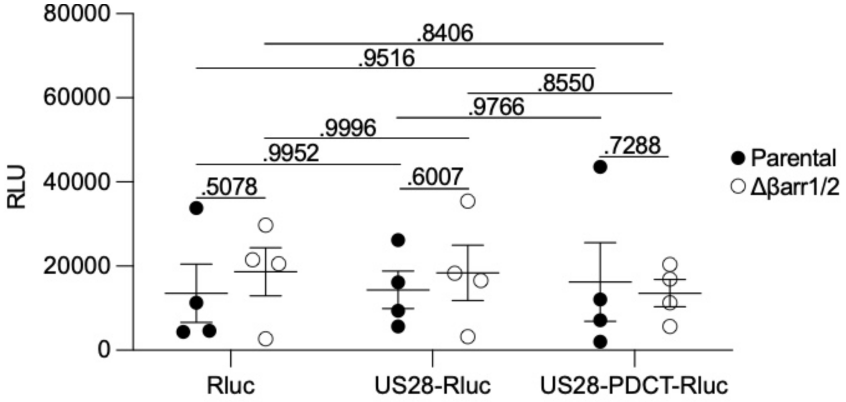


**Supplementary Fig. 2: Similar total expression of US28-Rluc, US28-PDCT-Rluc, or Rluc alone across parental and β-arrestin-deficient HEK-293 cells transfected transiently to evaluate the recruitment of mGq-GFP to the Rluc-fused constructs.** Data represent the luminescence expressed as relative luminescence units (RLU) monitored from parental and β-arrestin-deficient HEK-293 cells transiently transfected with US28-Rluc, US28-PDCT-Rluc or Rluc alone. Data are shown as means ± SEM of $n$ = 4 independent biological replicates performed in technical quadruplicates and were analyzed by two-way ANOVA with Tukey's post hoc test.

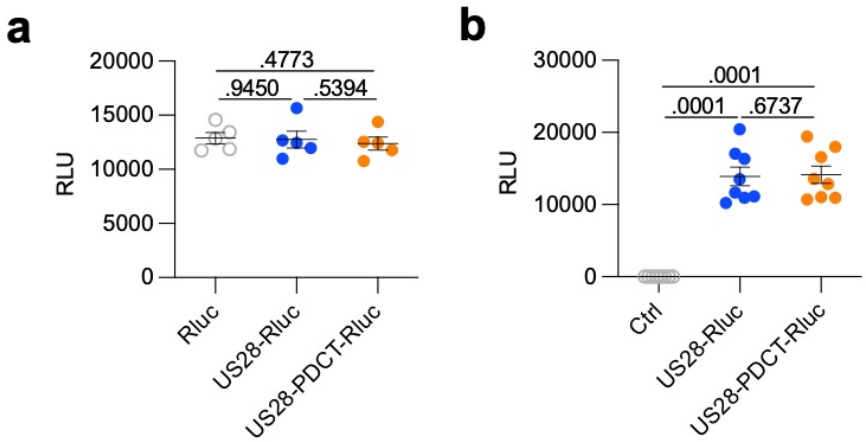


**Supplementary Fig. 3 : Similar total expression of US28-Rluc, US28-PDCT-Rluc, or Rluc alone across U251 cells transfected transiently to evaluate the constitutive production of second messengers. a** Luminescence expressed as relative luminescence units (RLU) monitored in U251 cells transiently transfected with US28-Rluc, US28-PDCT-Rluc, or Rluc alone used to monitor the constitutive DAG production. Data are shown as means ± SEM of $n$ = 5 independent biological replicates performed in technical quadruplicates. **b** RLU monitored from U251 cells transiently transfected with US28-Rluc, US28-PDCT-Rluc, or vector alone used to monitor the constitutive $IP_1$ production. Data are shown as means ± SEM of $n$ = 8 independent biological replicates performed in technical quadruplicates. Indicated $p$ values are derived from one-way ANOVA with Tukey's post hoc test.

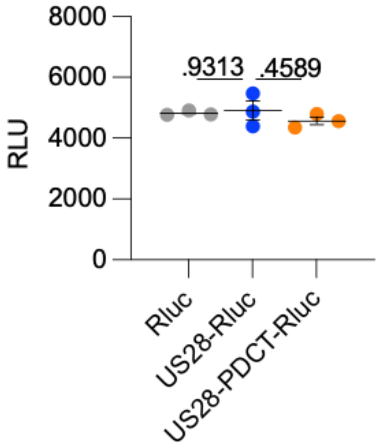


**Supplementary Fig. 4: Similar total expression of US28-Rluc, US28-PDCT-Rluc, or Rluc alone across U251 cells transfected transiently for RNA sequencing.** Luminescence expressed as relative luminescence units (RLU) monitored in U251 cells transiently transfected with US28-Rluc, US28-PDCT-Rluc, or Rluc used for RNA sequencing. Data are shown as means ± SEM of *n* = 3 independent biological replicates performed in technical triplicates or duplicates. One-way ANOVA with Dunnett's post hoc test.

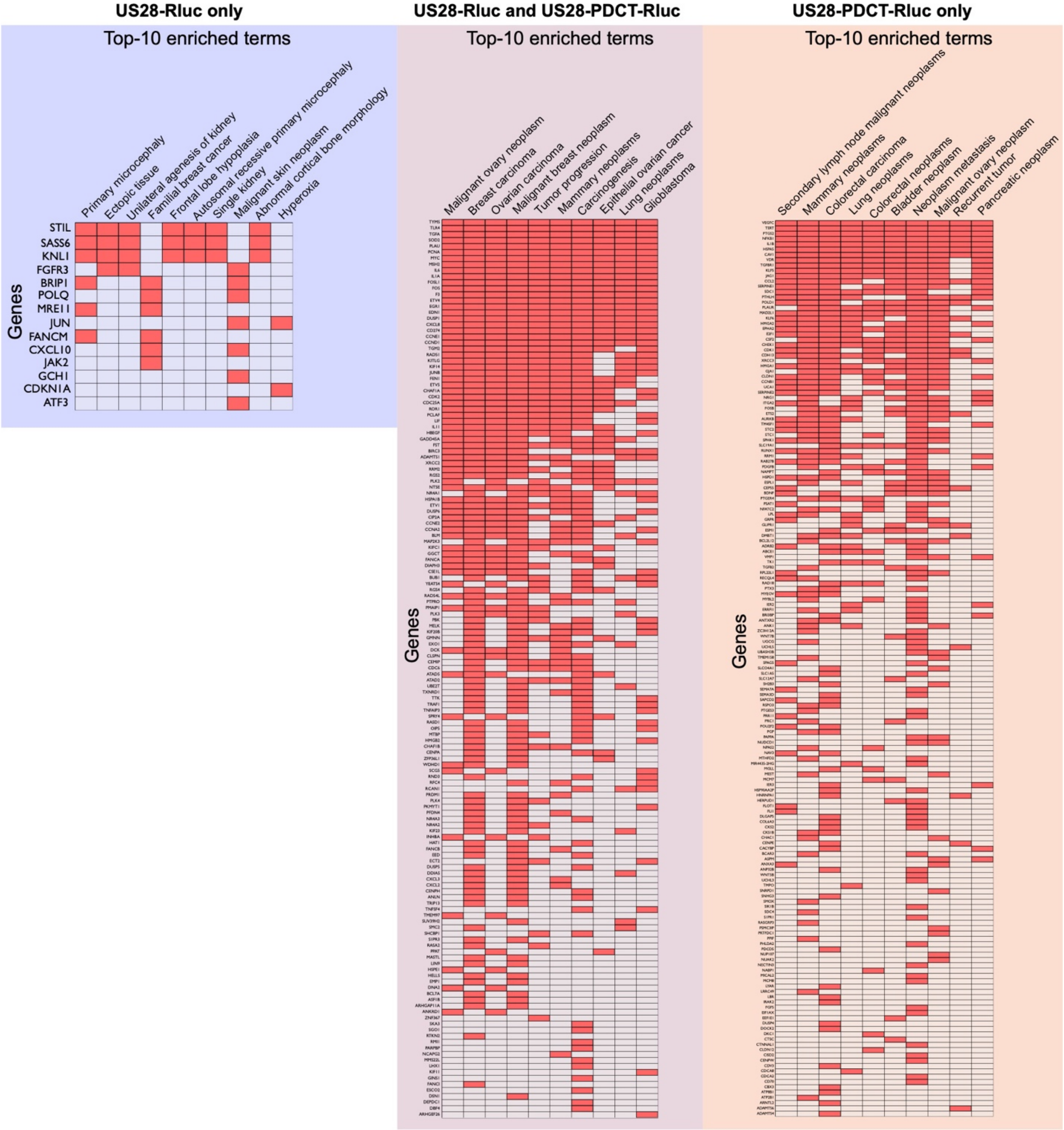


**Supplementary Fig. 5: Clustergram showing the top 10 enriched terms in the DisGeNET library with the associated input genes.** From the 83 genes only upregulated by US28-Rluc, the 247 genes upregulated by both US28-Rluc and US28-PDCT-Rluc, or the 347 genes upregulated by US28-PDCT-Rluc only.

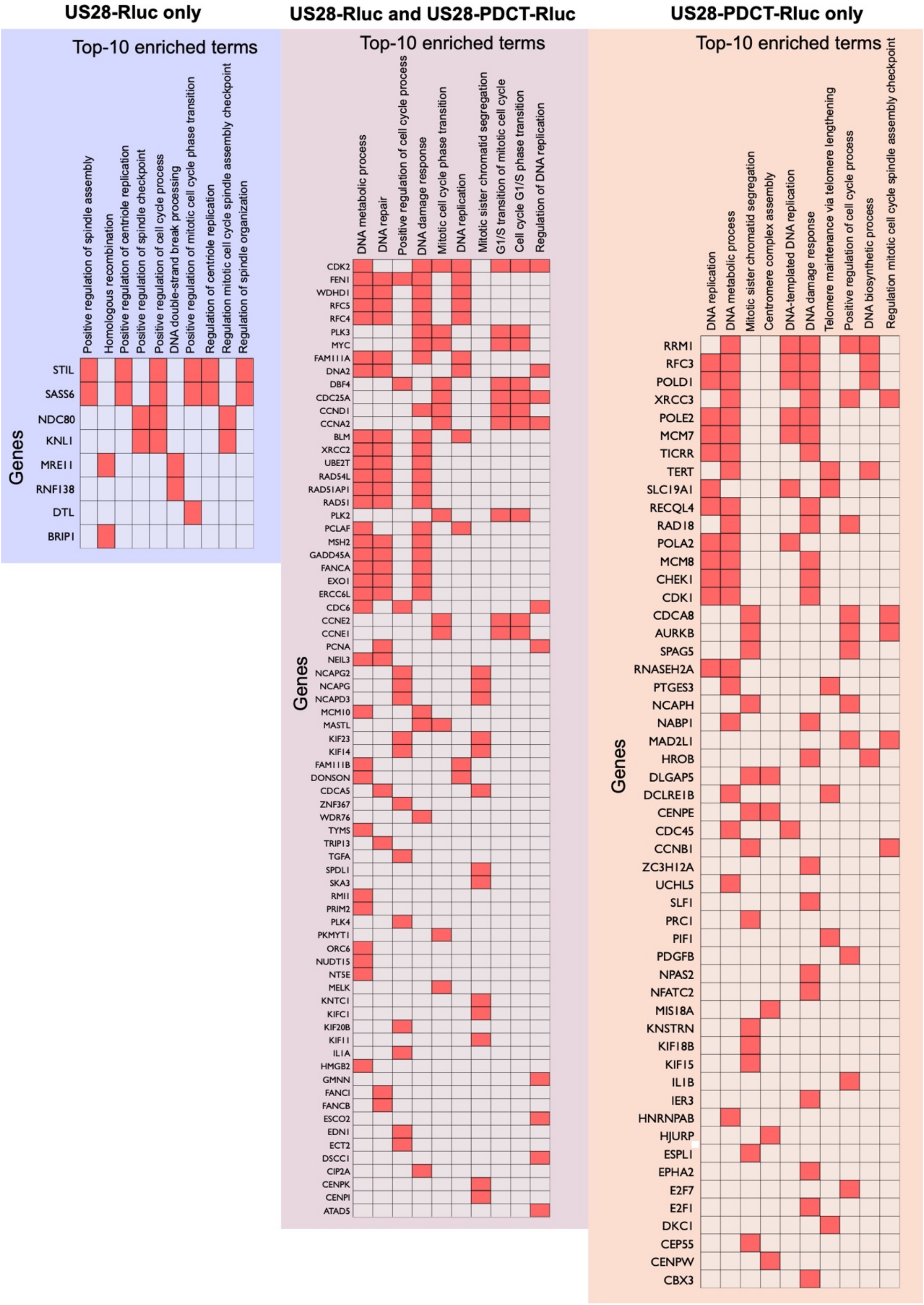


**Supplementary Fig. 6: Clustergram showing the top 10 enriched terms in the GO Biological Process 2025 library with the associated input genes.** From the 83 genes only upregulated by US28-Rluc, the 247 genes upregulated by both US28-Rluc and US28-PDCT-Rluc, or the 347 genes upregulated by US28-PDCT-Rluc only.

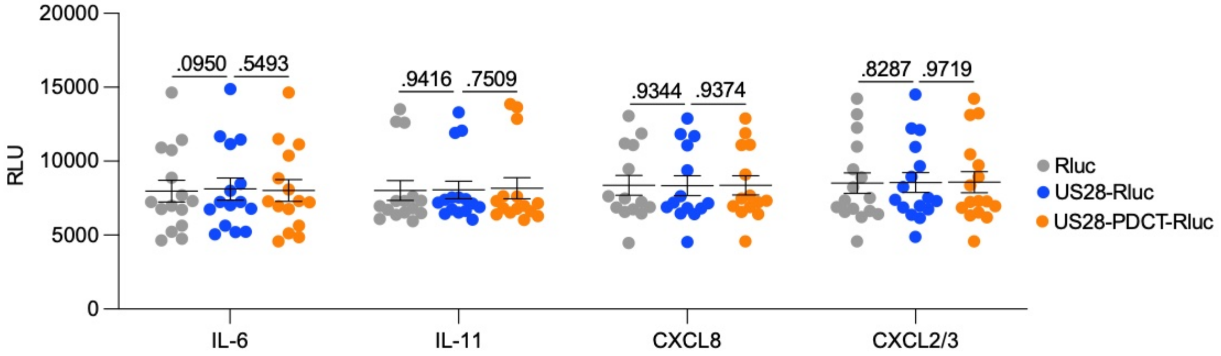


**Supplementary Fig. 7: Similar total expression of US28-Rluc, US28-PDCT-Rluc, or Rluc alone across U251 cells transfected transiently for cytokine secretion measurement.** RLU monitored in U251 cells transiently transfected with US28-Rluc, US28-PDCT-Rluc, or Rluc alone used to monitor the production of cytokines in the conditioned media. Data are shown as means ± SEM ($n$ = 15 for IL-6 and IL-11, $n$ = 14 for CXCL8, $n$ = 16 for CXCL2/3) performed in technical quadruplicates. Two-way ANOVA with Sidak's post hoc test.

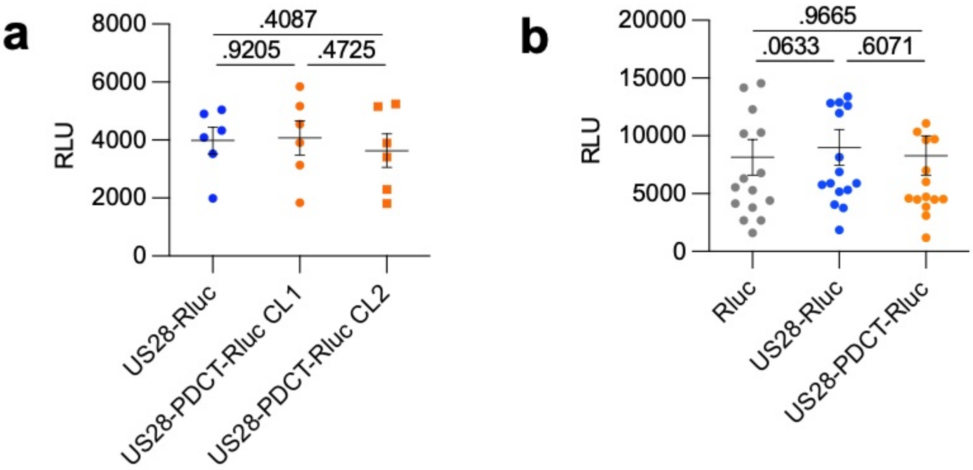


**Supplementary Fig. 8: Similar total expression of US28-Rluc, US28-PDCT-Rluc, or Rluc alone across U251 cells used for generation of spheroids or neutrophil migration assay. a** Luminescence expressed as relative luminescence units (RLU) monitored in clonal U251 cells stably expressing US28-Rluc, or US28-PDCT-Rluc used to generate spheroids. Data are shown as means ± SEM of $n$ = 6 independent biological replicates performed in technical quadruplicates. **b** RLU monitored from U251 cells transiently transfected with US28-Rluc, US28-PDCT-Rluc, or Rluc alone for which the conditioned media was used to monitor neutrophil migration. Data are shown as means ± SEM ($n$ = 16) performed in technical quadruplicates. **a,b:** one-way ANOVA with Tukey's post hoc test.

**Supplementary Table 1: RNA sequencing – raw data.** Raw data from FASTQ files from three independent biological replicates of RNA samples extracted from U251 cells expressing US28-Rluc, US28-PDCT-Rluc, or Rluc alone.

**Supplementary Table 2: Differential gene expression analysis (DGEA) – US28-Rluc versus Rluc.** DGEA from three independent biological replicates of RNA samples extracted from U251 cells expressing US28-Rluc or Rluc.

**Supplementary Table 3: Differential gene expression analysis (DGEA) – US28-PDCT-Rluc versus Rluc.** DGEA from three independent biological replicates of RNA samples extracted from U251 cells expressing US28-PDCT-Rluc or Rluc.

**Supplementary Table 4: List of genes upregulated by US28-Rluc only, US28-PDCT-Rluc only, or both.** From DGEA from three independent biological replicates of RNA samples extracted from U251 cells expressing US28-Rluc, US28-PDCT-Rluc, or Rluc.
